# scAmp analyzes focal gene amplifications at single-cell resolution

**DOI:** 10.64898/2026.02.14.705928

**Authors:** Matthew G. Jones, Natasha E. Weiser, King L. Hung, Xiaowei Yan, Sangya Agarwal, Jens Luebeck, Aditi Gnanasekar, Brooke E. Howitt, Ellis J. Curtis, Kevin Yu, John C. Rose, Katerina Kraft, Valeh Valiollah Pour Amiri, Leena Satpathy, Vineet Bafna, Paul S. Mischel, Howard Y. Chang

## Abstract

Oncogene amplification on extrachromosomal DNA (ecDNA) is a common driver of tumor progression and is associated with acquired drug resistance and poor patient survival. While whole genome sequencing (WGS) studies have revealed the landscape of genes amplified on ecDNA in tumors, it remains challenging to study the subclonal heterogeneity and functional (e.g., transcriptomic) consequences of ecDNA on tumors. To address this, we introduce *scAmp*: a probabilistic algorithm for detecting and analyzing ecDNA from single-cell datasets. We demonstrate *scAmp’s* improved accuracy over WGS approaches on well-characterized cell-lines and its applicability to clinical histopathology. We further showcase *scAmp* by analyzing 73 patient tumors profiled with single-cell ATAC-seq, where we analyze the subclonal evolution of ecDNA+ subclones and identify the effect of ecDNA amplifications on the chromatin accessibility landscape of cancer cells. Together, we anticipate that *scAmp* will broadly enable further studies – both retrospective and prospective – that dissect critical questions of how ecDNA affect cancer cells and the tumors in which they reside.

## MAIN

Focal amplification of DNA is a prevalent driver of tumor evolution^1,2^, but the mode of amplification can profoundly alter progression^3^. Circular extrachromosomal DNA (ecDNA, or “double-minutes”) amplifications are found in approximately 17% of primary tumors overall and are associated with significantly worse patient outcomes and increased drug resistance as compared to chromosomal amplifications^3–5^. While whole-genome sequencing (WGS) analyses of large patient cohorts have revealed the landscape of genes amplified on ecDNA^3,4,6–8^ (including canonical oncogenes like *MYC*, regulatory elements, and immunomodulatory genes like *SOCS1*), several questions remain that require increased resolution: for example, what is the distribution of ecDNAs among cancer cells within a tumor? How are ecDNAs associated with differential cancer cell programs and immune or stromal populations?

Current approaches for ecDNA detection focus on specific sequence features defined by WGS^8–10^. However, WGS-based approaches can have difficulty discriminating amplicons that persist as ecDNAs from those that have integrated back into chromosomes, often because the integrated amplicons retain the rearranged sequences and circular pattern of their ancestral ecDNAs. To overcome this limitation, we hypothesized that a defining feature of ecDNA biology – the non-Mendelian inheritance of individual ecDNA molecules during cell division^14^ – could enable an unbiased discovery of ecDNA from single-cell assays and address fundamental questions of how ecDNA affect tumor biology^9–11^ (**Figure 1**).

**Figure 1.**
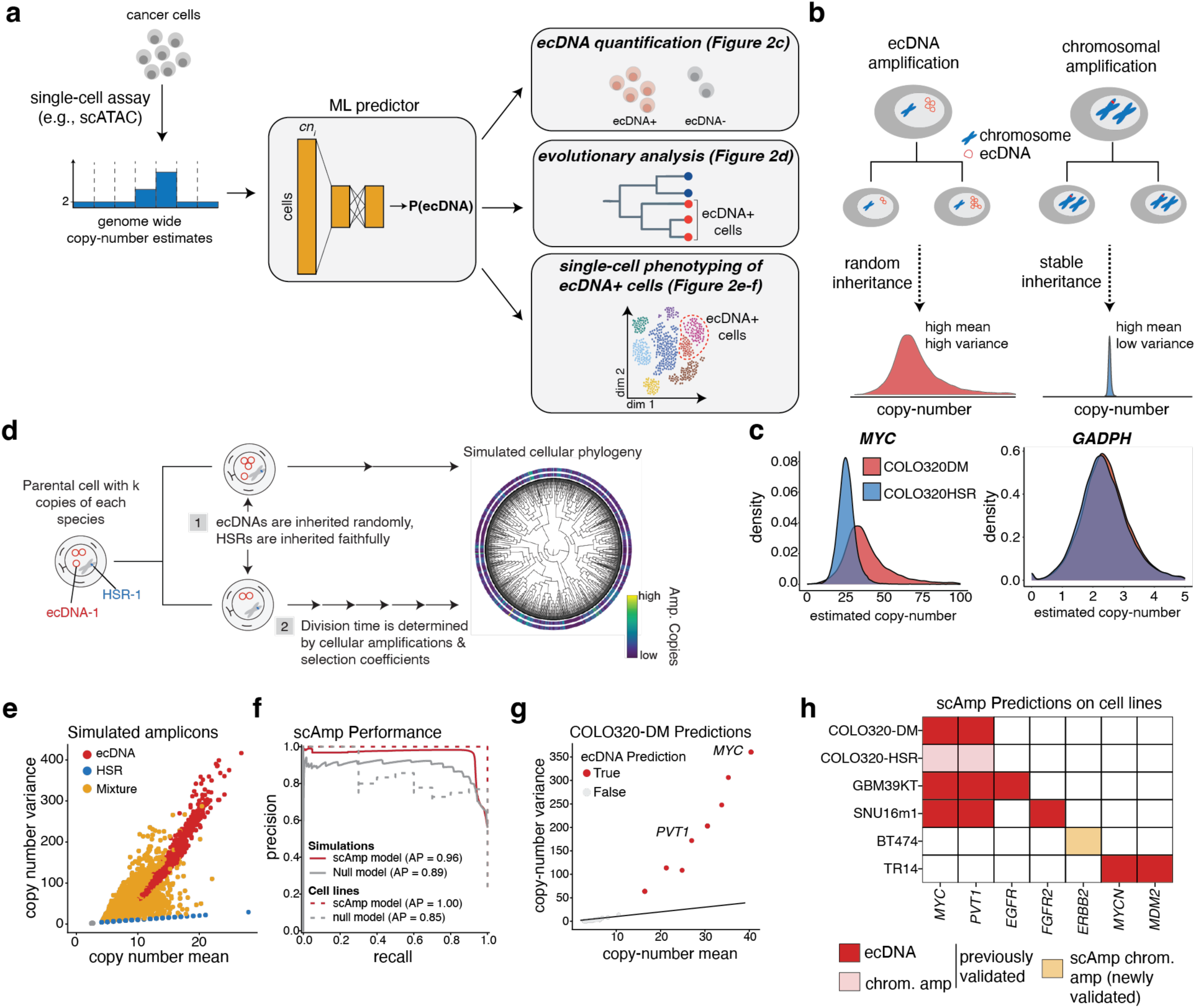
*scAmp* detects ecDNA-amplified genes from single-cell copy-number data. **(a)** An overview of the *scAmp* workflow. Copy numbers are inferred from individual cells (e.g., single-cell ATAC-seq data). For each gene, *scAmp* then infers the likelihood of that gene being amplified on ecDNA based on the properties of its copy-number distribution across cells. This enables downstream analyses such as quantifying the prevalence of ecDNA in a tumour, assessing an ecDNA’s evolutionary history, and investigating the phenotypic effects of ecDNA on cancer cell state. **(b)** A schematic representation of how random ecDNA segregation leads to greater copy-number variance. **(c)** Evaluation of single-cell copy-number distributions of *MYC* in COLO320-DM (ecDNA+) and COLO320-HSR (non-ecDNA), as compared to *GAPDH*. **(d)** A schematic representation of the simulation framework for generating training data for machine learning models. **(e)** Overview of simulated amplicon distributions from the simulation framework displayed in (d). **(f)** ROC curves showing performance of *scAmp* versus a “null” model based on mean copy-number alone on training data. **(g)** Summary of ecDNA predictions and copy-number distributions in the COLO320-DM cell line. Each point indicates an individual gene, and well-characterized genes (*MYC* and *PVT1*) are annotated. **(h)** Summary gene-level predictions from *scAmp* across cell lines; *scAmp* correctly predicts that *ERBB2* is chromosomally amplified, not extrachromosomally as inferred by WGS (**Extended Data Figure 5**).

Due to their lack of centromeres, ecDNA are randomly inherited during mitosis and thus drive greater copy-number variance in a population compared to chromosomal amplifications^14,15^ (**Figure 1b**). An analysis of the isogenic cell line pair COLO320-DM (containing ecDNA-amplified *MYC*) and COLO320-HSR (containing chromosomally-amplified *MYC*) underscores this hypothesis: while *MYC* is amplified in both cell lines, *MYC*’s copy-number variance is substantially greater in COLO320-DM cells (**Figure 1c**; **Extended Data Figure 1a**; **Methods**). Here, we leverage the discriminative ability of single-cell copy-number distributions to enable single-cell analysis of ecDNA with a new computational framework called *scAmp* (“single-cell amplicon” analysis; **Figure 1a**).

*scAmp* operates on single-cell copy-number data (which can be obtained from assays such as single-cell WGS^16,17^ or single-cell assay for transposase-accessible chromatin by sequencing [scATAC-seq]^18,19^) and predicts a likelihood for each gene that it is amplified on ecDNA. We trained models on simulated copy-number distributions of ecDNA and chromosomal amplification using a previously-described forward-time evolutionary model of focal amplifications^13,14^ and evaluated performance on held-out well-characterized cell lines^5,6,14^ and tumor samples from The Cancer Genome Atlas (TCGA) with paired WGS data^6,20^ (**Figure 1d; Extended Data Figure 1b***-***d; Methods**). We then selected the best-performing model by evaluating performance on held-out single-cell data (**Extended Data Figure 2; Methods**). From this analysis, we determined a best-performing multi-layer perceptron (MLP) with an average precision (AP) of 0.96 on simulated data (as compared to an AP of 0.89 for a logistic regression model trained just on mean copy-numbers; **Figure 1e-f**), perfect agreement with previous characterization of cell lines as opposed to baseline models (an AP of 1.0 for *scAmp* versus 0.85 for the null model; **Figure 1g-h; Extended Data Figure 3**; **Methods**), and an AUROC of 0.86 on predicting the ecDNA status of patient tumor samples (**Extended Data Figure 2** and **Extended Data Figure 4**). As expected, while the null model performs well overall, it cannot disambiguate between ecDNA and highly amplified chromosomal amplifications (e.g., average copy-number > 10) whereas *scAmp* remains accurate across copy-number regimes (**Extended Data Figure 2e-f** and **Extended Data Figure 3b**).

Cases in which *scAmp* disagreed with WGS revealed notable limitations of bulk genome sequencing for determining ecDNA status (**Figure 1f-h**). For example, the breast cancer cell line BT474 was predicted by WGS to have an ecDNA containing *ERBB2*^6^ while *scAmp* predicted *ERBB2* to be chromosomally amplified (**Figure 1h; Extended Data Figure 5**). Definitive characterization of *ERBB2’*s amplification mode by fluorescence *in situ* hybridization (FISH) of DNA on metaphase spreads of BT474 cells revealed that indeed *ERBB2* was chromosomally amplified, consistent with the prediction from *scAmp* (**Extended Data Figure 5; Methods**). The failure of WGS to correctly determine *ERBB2*’s amplification mode likely derives from either: (1) the ambiguity of head-to-tail reads that could resemble ecDNA or other amplifications like fold-back inversions (FBIs)^15^; or (2) the ancestral presence of ecDNA carrying *ERBB2* that has since integrated into the chromosome, retaining the discordant read structure that would support an ecDNA classification in bulk WGS analysis^6,14,21,22^. This result demonstrates that *scAmp* can predict the present status of ecDNA amplification with higher specificity than WGS approaches alone.

With *scAmp*’s ability to detect ecDNA status of genes at the single-cell level, we next turned to assessing its utility on patient samples and re-analyzed a recent cohort of 73 patient tumors profiled with scATAC-seq through The Cancer Genome Atlas^20^ (**Figure 2a; Methods**). As reported above, we found that *scAmp*’s predictions were largely concordant with WGS data alone (80% agreement on a per-gene level and 79% that a tumor contained at least one ecDNA-amplified gene; **Figure 2a**, **Extended Data Figure 4** and **Extended Data Figure 6a**). In this dataset, gliomas (GBMx), lung cancers (LUAD), and breast cancers (BRCA) had the highest prevalence of ecDNA (**Figure 2b**). Moreover, *scAmp* predicts key genes to be commonly amplified on ecDNA, including *EGFR*, *MYC,* and *KRAS* (**Extended Data Figure 6b**). Similar to the analysis of BT474, in the cases that *scAmp* disagrees with WGS, we find that the copy-number distribution supports *scAmp*’s predictions (**Extended Data Figure 6c-d**).

**Figure 2.**
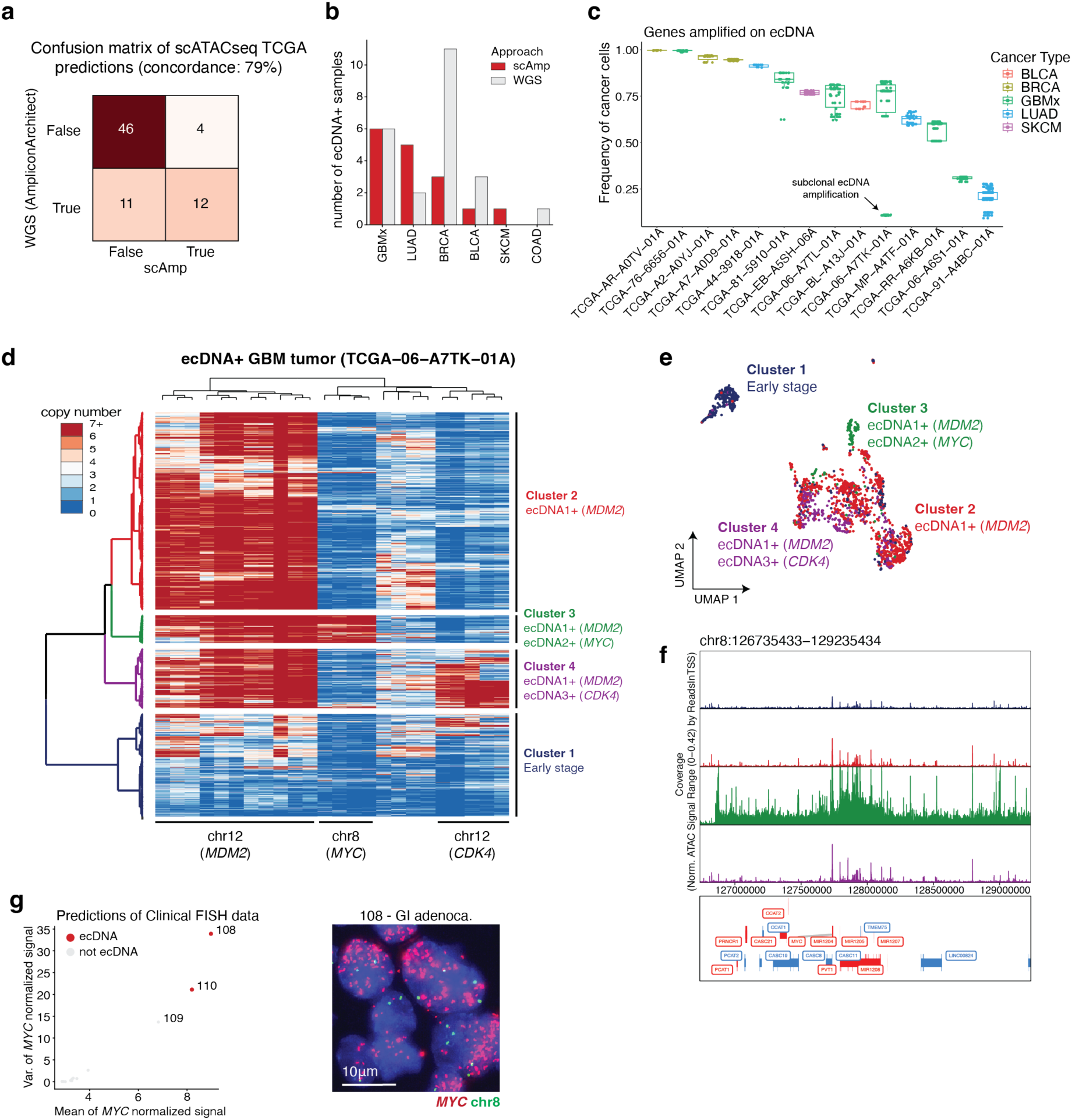
*scAmp* enables retrospective analysis and classification of patient tumours. **(a)** Confusion matrix reporting the concordance between *scAmp* and WGS predictions on a cohort of 73 patient tumours from TCGA. **(b)** Overview of ecDNA classifications in each tumour type, comparing *scAmp* and WGS calls. **(c)** Fraction of cancer cells with ecDNA amplifications across all ecDNA+ tumour samples. Each point is a gene predicted to be on ecDNA. Boxplots report the quartiles of the distributions, and whiskers extend to 1.5x the interquartile distance. **(d)** Clustered heatmap of ecDNA-amplified 3Mb windows in an example tumour sample, GBMx TCGA−06−A7TK−01A. Four hierarchical clusters are identified based on tree depth and annotated according to the representative oncogenes found on ecDNA amplifications (**Methods**). Chromosomes and representative focal genes are indicated below the heatmap. **(e)** UMAP projection of single-cell chromatin accessibility data, with colours corresponding to the hierarchical clusters presented in (d). **(f)** Chromatin accessibility tracks for a chromosome 8 region centred on *MYC*. Tracks are pseudo-bulked and coloured according to the hierarchical clusters identified in (d). **(g)** Predictions of ecDNA status of *MYC* amplifications from tissue microarray (TMA) data profiled with interphase DNA FISH (left). A representative interphase DNA FISH image of cancer cells from a gastrointestinal adenocarcinoma (“GI adeno.”) tumour predicted to be ecDNA+ (108; right). Scale bar indicates 10um.

Applying *scAmp* to assays like single-cell ATAC-seq also enables the phenotypic analysis of cell states within ecDNA+ tumors (**Extended Data Figure 7**). Stratifying by tumor type, we find substantial shifts in the immune composition of tumors: for example, T cells are significantly enriched in BRCA and LUAD tumors whereas macrophages were specifically increased in GBMx samples (**Extended Data Figure 7a-b**; **Methods**). We further leveraged the single-cell nature of this assay to stratify cancer cells within the same tumor by ecDNA status and inspect cell state changes (**Extended Data Figure 7c**). We found consistent changes in ecDNA+ cancer cells, including upregulation of glycolysis (*p*=0.02, Wilcoxon rank-sums test) and hypoxia-sensing pathways (*p*=0.06, Wilcoxon rank-sums test) as well as down-regulation of mitotic spindle assembly and reactive oxygen species signatures (*p*=0.12 and *p*=0.1, respectively, Wilcoxon rank-sums test).

The ability to detect ecDNA on a cell-by-cell basis provides the opportunity to infer the subclonal heterogeneity of ecDNA within tumors^11^. We find a striking variability in the fraction of ecDNA-amplified cancer cells across tumors: while ecDNA appear in almost every cell in some tumors, in others ecDNA amplifications may only be found in ∼10% of cancer cells (**Figure 2c**). The distribution of ecDNA within a single tumor may provide insight into the relative evolutionary timing of ecDNA biogenesis. For example, in one GBMx tumor (TCGA−06−A7TK−01A) we find that while ∼80% of cells contain an ecDNA amplifying *MDM2*, two distinct subsets of cells also contain an additional ecDNA: one encoding *MYC* (Cluster 3) and another encoding *CDK4* (Cluster 4) (**Figure 2d**). These data suggest that the additional ecDNAs may have originated in a subclone of the larger ecDNA+ population, either in response to ongoing genomic instability or selection pressure imparted from the tumor microenvironment^23,24^. We also find that regulatory changes in chromatin accessibility accompany these changes in ecDNA content of subpopulations: cells with the subclonal *MYC* ecDNA cluster separately and exhibit highly accessible chromatin of the amplified chromosome 8 locus as compared to the cells that do not amplify this *MYC* ecDNA, consistent with previous findings that ecDNA chromatin is highly accessible^25^ (**Figure 2e-f**).

Finally, we tested if *scAmp* generalizes beyond single-cell genomic assays to enable the classification of ecDNA+ tumors from clinically relevant modalities like DNA FISH in formalin-fixed paraffin embedded (FFPE) tissues. To establish the utility of *scAmp* for ecDNA calling in FFPE, we first confirmed that *scAmp* could accurately distinguish ecDNA amplifications from chromosomal amplifications in FFPE tissue from mouse xenografts tumors of COLO320-DM and COLO320-HSR cells (**Extended Data Figure 8a; Methods**). We next extended these results to demonstrate the feasibility of this approach on patient tissues with a proof-of-concept analysis of a tissue microarray (TMA) generated from 14 tumors reported to have focal amplifications in the Stanford Pathology archive, including five tumors reported to amplify *MYC* (**Figure 2g; Extended Data Figure 8b**). In the ∼1mm tissue cores represented on the TMA, we detected *MYC* ecDNA amplifications in 2 samples and *MYC* chromosomal amplification in 1 sample, consistent with the overall appearance of FISH signal in these tumors (**Figure 2g**; **Extended Data Figure 8c**; **Methods**). Together, these analyses demonstrate *scAmp*’s utility in a range of contexts, including basic studies of tumor biology and potentially in clinical diagnosis.

## ACKNOWLEDGEMENTS

We thank Anton Henssen, Andrea Ventura, and all members of the Chang and Mischel laboratories for helpful discussions. This work was delivered as part of the eDyNAmiC team supported by the Cancer Grand Challenges partnership funded by Cancer Research UK (CGCATF-2021/100012 (P.S.M. and H.Y.C.) and CGCATF-2021/100025 (V.B.)) and the National Cancer Institute (OT2CA278688 (P.S.M. and H.Y.C.) and OT2CA278635 (V.B.)). M.G.J. is supported by NIH K99-CA286968. N.E.W. is supported by NIH K08-CA296931. X.Y. was supported by Damon Runyon Cancer Research Fellowship. K.L.H. was supported by a Stanford Graduate Fellowship and an NCI Predoctoral to Postdoctoral Fellow Transition Award (NIH F99CA274692). V.V.P.A. is supported by a Stanford HAI Graduate fellowship and the Amazon Core AI fellowship.

H.Y.C. was an Investigator of the Howard Hughes Medical Institute. B.E.W. is supported by the Irene Adler Endowed Scholar fund. Research reported in this publication was supported by the National Cancer Institute under Award Number P30CA124435 (B.E.H.). The content is solely the responsibility of the authors and does not necessarily represent the official views of the National Cancer Institute.

## AUTHOR CONTRIBUTIONS

H.Y.C and M.G.J. conceived the project with insights from J.C.R., J.L., P.S.M., and N.E.W.

M.G.J. developed the *scAmp* algorithm and performed benchmarking. K.L.H. and M.G.J. generated scATAC data and analysed it, with support from V.V.P.A. and L.S. S.A. and X.Y. cultured cells and performed imaging of the BT474 cell line with support from K.K.. J.L. provided annotations of patient samples from TCGA with supervision from V.B. A.G. and E.J. performed xenograft experiments of COLO320-DM and COLO320-HSR cell lines and prepared slides for DNA FISH imaging by N.E.W. B.H. and N.E.W. prepared a TMA of cancer samples and N.E.W. performed DNA FISH imaging and quantification. M.G.J. deployed *scAmp* on all samples described in this study with support from L.S. and V.V.P.A.

M.G.J. wrote the *scAmp* software package with support from K.Y. M.G.J., N.E.W. and H.Y.C. wrote the manuscript with input from all authors.

## DATA AND CODE AVAILABILITY

The *scAmp* software is publicly available on GitHub at https://github.com/mattjones315/scamp. New sequencing data and source imaging data for this study will be deposited and made available upon publication. AmpliconClassifier outputs containing ecDNA coordinates in TCGA samples were previously published and are publicly available at figshare (https://doi.org/10.6084/m9.figshare.24768555.v1).

Previously published scATAC-seq data of cell lines are publicly available at (GSE159986, SRS21730888, SRS21730887 SRS21730902, and SRS21730907). Previously published TCGA scATAC-seq data are publicly available via NCI Genomic Data Commons (https://gdc.cancer.gov/about-data/publications/TCGA-ATAC-Seq-2024).

## METHODS

### Generating simulated training data

To generate data for training models, we simulated trajectories of genes that were amplified on either ecDNA or chromosomes (“HSRs”) using a modified Cassiopeia simulation framework^26^ (see section “**Simulation of ecDNA and HSR copy-number distributions”**). We simulated 1,000 trajectories for both 1M and 500k cells (a total of 2,000 trajectories): for each trajectory, we obtained copy-number for an ecDNA amplicon and an HSR amplicon for each cell (copy-numbers are denoted as 𝑐_j_^a^ for cell *i* and amplicon *a*). Then, we created 10,000 training datasets by sampling a random trajectory of ecDNA and HSR amplicons and applying a noise model to the observed copy-numbers per cell to account for noise in the sampling and copy-number estimation process. The noise model consisted of simulating an under-dispersed normal distribution parameterized by the observed counts: specifically, given an observed copy-number 𝑐_j_^a^ for cell *i* and amplicon *a*, we drew modified counts from the following distribution:

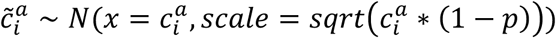

where 𝑝 ∼ 𝑈𝑛𝑖𝑓(0.1, 1). Importantly, we elected to simulate under-dispersed copy-numbers by inspecting the distributions of HSR-amplified genes in real single-cell data.

We additionally simulated 500 cell populations without any amplification by drawing copy-numbers from a mixture model as follows:

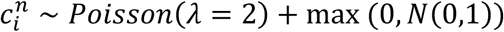

Finally, we simulated two mixtures of ecDNA/HSR and amplified/non-amplified cell populations: first, we simulated 10,000 mixtures of cells that amplified both ecDNA and HSR genes by merging copy-number distributions for both ecDNA and HSR and a random proportion. Second, we simulated 1,000 mixtures of non-amplified cells and cells that either amplified ecDNA or HSR genes. The same noise model was applied to copy-numbers for these mixtures as above. Altogether, this resulted in 32,500 training examples: 10,000 ecDNA distributions; 10,000 HSR distributions; 10,000 ecDNA/HSR distributions; 1,000 normal/ecDNA distributions; 1,000 normal/HSR distributions; and 500 normal distributions.

### Simulation of ecDNA and HSR copy-number distributions

We used a modified evolutionary model of ecDNA copy-number dynamics^13,14^ to simulate trajectories of both ecDNA and HSR amplifications. Simulations began with a single cell with 𝑘^e^ ecDNA and 𝑘^c^chromosomal copies of a given gene, as determined below:

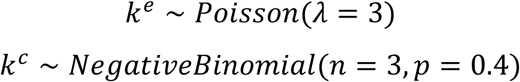

Simulations were additionally parameterized with selection coefficients acting on both ecDNA+ and HSR+ cells (𝑠_e_, 𝑠_c_, respectively); a base birth rate (𝜆_base_ = 0.5); and a death rate (𝜇 = 3).

Evolutionary simulations then proceeded as previously described^13,14^: starting with the parental cell, a birth rate is defined based on the selection coefficient acting on the cell (a combination of 𝑠_e_ and 𝑠_c_, depending on whether the cell has ecDNA and/or chromosomal copy amplifications) as 𝜆_1_ = 𝜆_base_ × (1 + 𝑠^’^) where 𝑠^)^is the combination of selection coefficients from both ecDNA and HSR. Then, a waiting time until the next cell division is drawn from an exponential distribution, parametrized by the updated birth rate:

𝑡_b_ ∼ exp (−1/𝜆_1_). Time to cell death is also drawn from an exponential distribution: 𝑡_d_ ∼ exp (−𝜇). When 𝑡_b_ < 𝑡_d_, a cell division event is simulated and a new edge is added to the growing phylogeny with edge length 𝑡_b_; otherwise, the cell dies and the lineage is stopped. This process will continue until a user-defined stopping condition is specified: in this case, the number of extant (living) cells being either 1M or 500k.

During a cell division, ecDNAs are split among daughter cells randomly, as previously described and validated^14^, generating new daughter cell ecDNA counts of 𝑘_d1_^e^ and 𝑘_d2_^e^ for 𝑑_1_ and 𝑑_2_. Specifically,

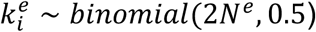

Where 𝑁^e^ is the ecDNA copy-number of the parent. Conversely, the chromosomal copy number of the daughter cells is kept constant, assuming faithful chromosome inheritance^15^.

### Creating ground-truth cell line annotations

We created validation datasets by first curating data from previously characterized cell lines. We elected to focus on 6 cell lines that were known to contain either ecDNA or chromosomal amplifications, per previous characterization by a combination of WGS and/or DNA FISH on metaphase spreads (as detailed in **Extended Data Figure 1b**): COLO320-DM, COLO320-HSR, GBM39-KT, SNU16 (and SNU16m1), TR14, and BT474.

While several genes are amplified on a given ecDNA or HSR, we elected to evaluate a model’s performance primarily on its ability to successfully predict the status of a well-characterized genes, including *MYC*, *PVT1, EGFR, FGFR2, ERBB2, MYCN*, and *MDM2*.

### Creating ground-truth TCGA annotations

We created ground-truth annotations of gene amplification status from WGS data sequenced through The Cancer Genome Atlas (TCGA). Specifically, we utilized results from the AmpliconSuite^6,10^ pipeline, which detects genomic amplifications and classifies them as ecDNA or other forms of amplifications (e.g., complex non-cyclic or linear); these results were previously published^3,6,14^. Briefly, this approach for detecting ecDNA uses three steps wrapped in the AmpliconSuite pipeline (https://github.com/AmpliconSuite/AmpliconSuite-pipeline, v1.1.1). First, given a BAM file, the analysis pipeline detects “seed” intervals where copy-number amplifications exist (CN > 4.5, with a size between 10kb-10Mb). Copy-numbers are pre-computed from the BAM file using a workflow such as cnvkit^27^. Second, AmpliconSuite invokes AmpliconArchitect^10^ (AA) to perform joint copy-number and breakpoint detection analysis in the focally-amplified regions, forming a copy-number-weighted breakpoint graph. AA then extracts paths from this breakpoint graph, each of which corresponds to a particular amplification, that collectively explain the copy-number and breakpoint patterns detected. Finally, a rule-based classification is performed using AmpliconClassifier^23^ based on the paths extracted by AA to predict the mode of focal amplification (ecDNA or other). This includes assessing structural variant types, copy-numbers, and structure of the path (e.g., if the path is circular or not). The complete classification criteria and description of the AmpliconClassifier tool are available in the supplementary information of ref^11,24,25^.

From the AmpliconSuite analysis, we created two annotations that we used for model evaluation: first, we annotated tumours by whether or not they had any ecDNA amplification (sample-level); and second, within each tumour we annotated which genes were predicted to be amplified on ecDNA based on AmpliconSuite analysis (gene-level). We evaluated model performance on both criteria, as reported in **Extended Data Figure 4.**

### Training and benchmarking *scAmp*

We built models that gene amplifications as either ecDNA+ or non-ecDNA based on a featurization of single-cell copy-number distributions. While in principle the copy-number distribution could be featurized in many ways, in this proof-of-concept work we elected to reduce this distribution to simple statistics. Specifically, given a distribution of copy-numbers of cells, 𝐶 = {𝑐_1_,…, 𝑐_n_}, we extracted 14 features (𝑋): the mean, variance, coefficient of variation (variance / mean), the deciles of the copy-number distribution, and the inter-quartile range. While computing these statistics, we only considered “amplified” cells in which the copy-number exceeded 2.

Using these features, we trained fully-connected neural networks (Multilayer Perceptron [MLPs]) using pytorch^28^ (v2.6.0.dev20241219). We performed hyperparameter searches across the following parameters (**Extended Data Figure 2**):

- Number of layers: [0, 1, 2, 3, 4] (0 corresponding to a basic logistic regression)
- Size of hidden layers: [10, 20, 30]
- Learning rates: [1e-3, 1e-4]
- Activation functions: [“relu“, “tanh”, “sigmoid”]
- Batch size: [64, 256, 4096]

We trained models across 400 epochs using the simulated copy-number data (see above “**Generating simulated training data**”) and assessed performance on held-out single-cell data from cell lines and patient tumours (see above “**Creating ground-truth cell line annotations”** and “**Creating ground-truth TCGA annotations”**, respectively). For the purposes of this supervised learning task, we classified any mixed HSR/ecDNA sample that had at least 10% ecDNA+ cells as ecDNA+, and otherwise non-ecDNA. We trained 5 models each with different seeds to assess the robustness of the model. We prioritized models based on both the per-sample AUROC of and accuracy of predictions on patient tumour data (i.e., predicting whether or not a patient had *any* gene amplified on ecDNA). The final *scAmp* model (v1.0.0) consisted of 3 hidden layers with 20 neurons each and a “sigmoid” activation function (nn.Sigmoid). While training, we used a learning rate of 1e-4, a batch size of 64, and a dropout rate of 0.1 to reduce overfitting.

We also benchmarked *scAmp* against a RandomForest classifier (trained with sklearn version 1.6.0 using default parameters) and a “null” linear model trained on just the mean copy-number. We report the performance of these models on held-out cell line data **Extended Data Figure 3** and compare the performance of RandomForest to *scAmp* on held-out tumour data in **Extended Data Figure 4**.

### Cell culture

BT-474 cell line was obtained from American Type Culture Collection (ATCC) [Cat# CRL-3247] and was cultured in standard culture media (RPMI 1640 + 10% FBS, 1% penicillin/streptomycin (50IU/ml), 1% HEPES, 1% Glutamax) (Gibco, Life Technologies) at 37°C in 5% CO_2_.

### Metaphase DNA FISH of BT474

Mitotic cells were arrested with 0.1 µg/mL colcemid for 4 h, collected by centrifugation, and washed once with phosphate-buffered saline (PBS). The cell pellet was then incubated in 0.075 M KCl for 20 min at 37 °C to induce hypotonic swelling. Cells were subsequently fixed and washed three times with freshly prepared fixative consisting of methanol and glacial acetic acid (3:1, v/v). Fixed cells were resuspended in an appropriate volume of fixative and dropped from a height of several inches onto pre-humidified glass slides to achieve optimal chromosome spreading.

Slides were rinsed with PBS, washed in 2× saline-sodium citrate (SSC), and sequentially dehydrated in 70%, 85%, and 100% ethanol. Hybridization probe solution was applied, and the slides were denatured at 75 °C for 3 min followed by hybridization overnight at 37 °C. Post-hybridization washes were performed in 2× SSC containing 0.1% Tween-20 and PBS. Slides were then mounted with ProLong™ Gold Antifade Mountant containing DAPI.

### Paired scATAC-seq and scRNA-seq library generation

Single-cell paired RNA-seq and ATAC-seq libraries were generated with the 10X Chromium Single-Cell Multiome ATAC + Gene Expression kit according to the manufacturer’s protocol. Libraries were sequenced on an Illumina NovaSeq 6000 system. Single cell Multiome data for other cell lines were obtained from the following public databases: COLO320-DM & COLO320-HSR (GEO accession GSE159986), GBM39-KT (SRA accension SRS21730888), SNU16 (SRA accension SRS21730887), SNU16m1 (SRA accension SRS21730902), and TR14 (SRA accension SRS21730907).

### Copy-number analysis of scATAC-seq data

We calculated copy-numbers in 3Mb windows based on the background ATAC-seq signal as we previously described and validated^14,18,29^. Briefly, we determined read counts in large intervals across the genome using a sliding window of 3Mb moving in 1Mb increments across the reference genome (hg19). Genomic regions with known mapping artifacts were filtered out using the ENCODE hg19 blacklist. For each interval, insertions per bp were calculated and compared to 200 of its nearest neighbours with matched GC nucleotide content. The mean log_2_[fold change] was computed for each interval. Based on a diploid genome, copy numbers were calculated using the formula 𝐶𝑁 = 2 × 2^log2[FC]^ where CN denotes copy number and FC denotes mean fold change compared to neighbouring intervals. To query the copy numbers of a gene, we obtained all genomic intervals that overlapped with the annotated gene sequence and computed the mean copy numbers of those intervals.

### Analysis of scATAC-seq data from cell lines

Single-cell Multiome data was aligned to hg19 using cellranger-arc count (10X Genomics, v. 1.0.0). For the purposes of this study, we focused on the scATAC-seq fraction of the data and performed analysis using ArchR^30^ (v. 1.0.2). Cells with fewer than 1000 unique fragments or TSS enrichment less than 4 were filtered out. Doublets were removed using ArchR’s *addDoubletScores* and *filterDoublets* commands with the following parameters: k=10, knnMethod=“UMAP”, LSIMethod=1. Dimensionality reduction was performed using Iterative Latent Semantic Indexing with the *addIterativeLSI* function. For each cell line, we converted single-cell copy-number distributions of amplified genes (those with average copy-number exceeding 2.5) to *scAmp*-compatible features with the *prepare_copy_numbers* function with the following parameters: min_copy_number=3, max_percentile=99.9. Genes with predicted likelihood exceeding 0.6 were marked as ecDNA+.

### Predicting gene amplification status from TCGA scATAC-seq data

Single-cell ATAC-seq data for patients from TCGA was obtained from NCI Genomic Data Commons (https://gdc.cancer.gov/about-data/publications/TCGA-ATAC-Seq-2024) and copy-numbers were computed as described above (see section entitled “**Copy-number analysis of scATAC-seq data”).** For each patient, we converted single-cell copy-number distributions of amplified genes (those with average copy-number exceeding 2.5) to *scAmp*-compatible features with the *prepare_copy_numbers* function with the following parameters: min_copy_number=3, max_percentile=99.9. Genes with predicted likelihood exceeding 0.6 were marked as ecDNA+.

To quantify the frequency of ecDNA+ cells in **Figure 2c**, we computed the fraction of cells that amplified a gene predicted to be on ecDNA at 4 copies or more. To assess the phenotypic consequences of ecDNA on cancer cells, we first computed module scores for each signature in the MSigDB^31^ Hallmark set (available at https://www.gsea-msigdb.org/gsea/msigdb/human/genesets.jsp?collection=H) with ArchR (version v. 1.0.2) using the *addModuleScore* function with useMatrix=’GeneScoreMatrix’. We then performed *in silico* separation of cancer cells based on whether they amplified any gene predicted to be on ecDNA at 4 copies or more and assessed differential signature scores based on this *in silico* separation.

We also analysed the shifts in immune and stromal composition based on the ecDNA status of a given tumour (**Extended Data Figure 7a-b**). To do this, we separated immune and stromal populations from epithelial populations and annotated cells as previously described^20^.

### Analysis of GBMx sample TCGA−06−A7TK−01A

To analyse the subclonal evolution of the GBMx sample TCGA−06−A7TK−01A, we first inspected the shared patterns of copy-number amplifications in cancer cells computed across 3Mb windows (see above “**Copy-number analysis of scATAC-seq data**”). To identify regions that were likely ecDNA+, we focused on 3Mb regions that were amplified at greater than 7 copies in at least 5% of cells. We then performed hierarchical clustering of cells based on these regions using Euclidean distance and “ward” linkage and extracted 4 clusters based on tree depth.

To create the single-cell UMAP projection of this patient’s data, we performed analysis with ArchR (version 1.0.2). We first performed dimensionality reduction using Iterative Latent Semantic Indexing with the *addIterativeLSI* function with the following parameters: useMatrix=‘TileMatrix’, iterations=2, clusterParams=list(resolution=c(0.2, 0.4), n.start=10), and varFeatures=30000). We computed a UMAP projection from the LSI dimensions using nNeighbors=15, minDist=0.2, and metric=‘cosine’. Genome tracks were visualized using the *plotBrowserTrack* function using tileSize=100, upstream=1000000, and downstream = 1500000.

### Xenografts

Female athymic nude mice (Charles River Laboratories) were housed under standard conditions. Mice were anesthetized using isoflurane in an induction chamber immediately before subcutaneous cancer cell injection. 500,000 cells suspended in 100 µL of a 1:1 PBS:Matrigel mixture were injected subcutaneously into each flank bilaterally using a 25-gauge needle and a 1 mL syringe. Mice were monitored for signs of distress or complications after injection. Tumor length and width were measured several times per week until Day 21, at which time mice were sacrificed using a sealed CO₂ chamber, with cervical dislocation performed as a secondary confirmatory method. Tumors were sharply dissected from the subcutaneous space, immediately flash-frozen in liquid nitrogen, and stored at-80 °C. Excised tumor tissues were sectioned onto glass slides by the Stanford Human Pathology/Histology Service Center.

### DNA FISH on FFPE

Tumors with amplified oncogenes were identified from the Stanford Pathology archive in accordance with IRB protocol #69198 which was approved with the Stanford University Institutional Review Board. DNA FISH was performed using the Agilent Technologies histology FISH accessory kit (K579911-5) according to manufacturer’s instructions. Briefly, slides were incubated at 60°C for 1 hr prior to deparafinization in histoclear (2×5 min) and rehydration (2x 2min in 100% ethanol, 2x 2min in 70% ethanol). Samples were incubated in pre-treatment solution in a vegetable steamer for 20 minutes and then cooled at room temperature for 15 minutes. Samples were washed twice in wash buffer for 3 minutes. Samples were then treated with ready-to-use pepsin drops for 20min at room temperature followed by 2x 3min wash buffer incubations. Samples were dehydrated with a graded series of ethanol: 2 min each 70%, 85%, 100% ethanol. FISH probes targeting *MYC* (RP11-440N18) and the chromosome 8 centromere (CHR08-20-GR) with hybridization buffer were purchased from Empire Genomics After application of FISH probes diluted in hybridization buffer, slides were incubated at 82°C for 5min and then overnight at 37°C in a humid chamber. Samples were then incubated in stringent wash buffer at 65C for 10min, and then in wash buffer for 2×3min at room temperature. Slides were dehydrated in a series of ethanol washes: 2min each in 70%, 85%, and 100% ethanol. After air-drying, samples were mounded with the provided fluorescence mounting medium. Images were acquired on a Leica DMi8 widefield microscope using a 63x oil objective. Z-stack images were post-processed using Small Volume Computational Clearing on the LAS X thunder imager prior to generating maximum intensity projections. Image analysis was performed using the Aivia 15 (patient tumors) and Aivia 13 (xenograft) software. Green and red pixel classifiers were used to distinguish centromere 8 and *MYC* signals respectively. Segmentation was performed using the Cellpose module in Aivia 15 (patient tumors) and Aivia 13 (xenograft). Cells with circularity <0.4 or nuclear area >150µm were presumed to be poorly segmented and were filtered out prior to copy number analysis. Normalized *MYC* signal was computed by dividing the number of *MYC* foci by the number of chromosome 8 foci, and passed into *scAmp* for ecDNA classification.

## EXTENDED FIGURES

**Extended Data Figure 1.**
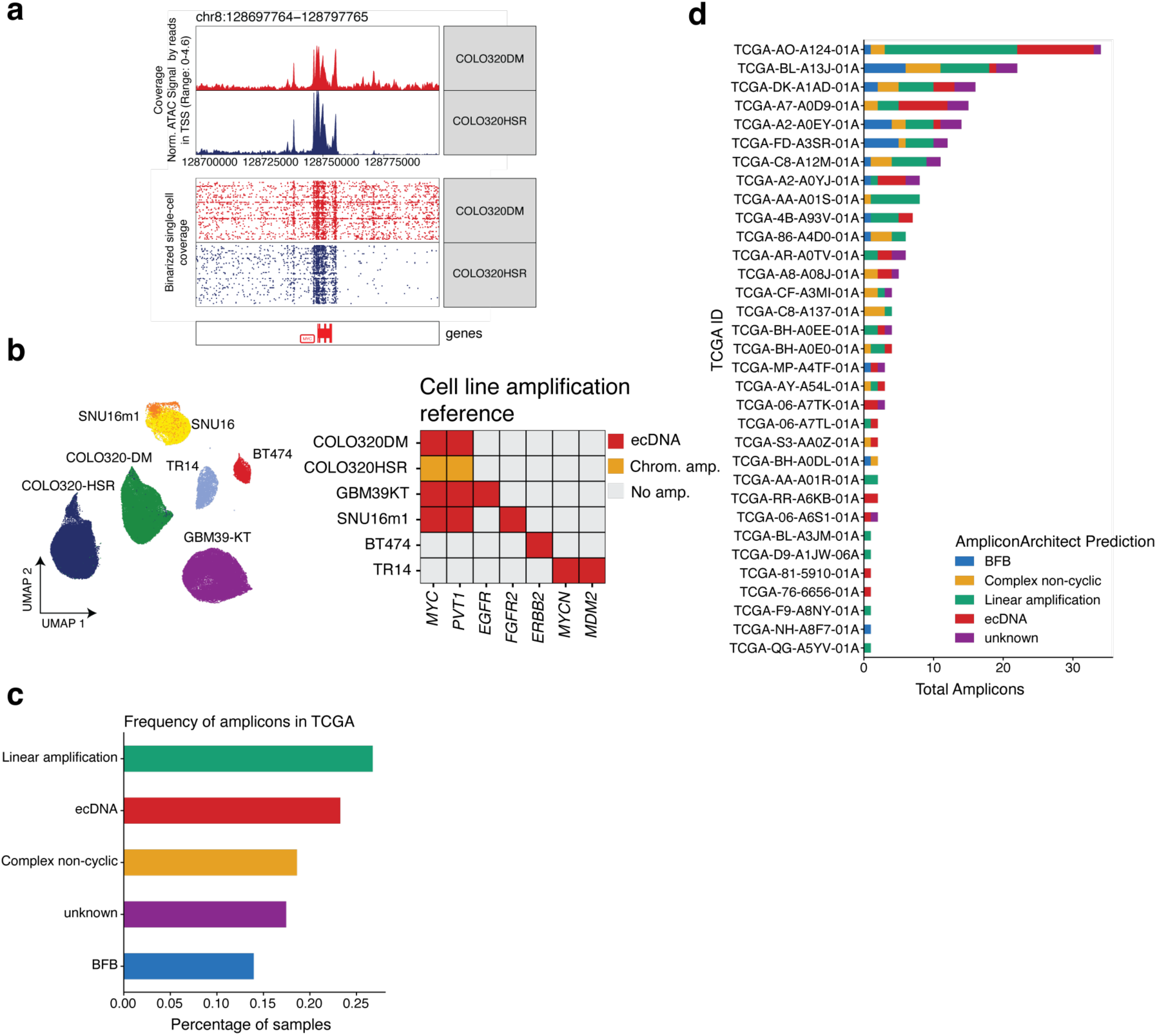
Overview of cell line and patient tumour annotations. **(a)** Single-cell ATAC-seq coverage for COLO320DM and COLO320HSR cell lines in the *MYC* locus. Pseubulk coverage (top) and binarized coverage of individual cells (bottom) demonstrate that while the average signal is similar, the COLO320DM ATAC signal appears to have greater variability. **(b)** A UMAP projection of single-cell ATAC-seq data for cell lines used int his study (left) and a summary of the characterized amplification mode of specific genes (right). Characterization of amplifications is derived from prior reports. **(c)** Summary of gene amplification annotations in patient tumours profiled with scATAC-seq from TCGA, as obtained from WGS analysis with AmpliconSuite. **(d)** The composition of amplicons per patient sample profiled with scATAC-seq from TCGA, where amplicon type is inferred from AmpliconSuite analysis of WGS data.

**Extended Data Figure 2.**
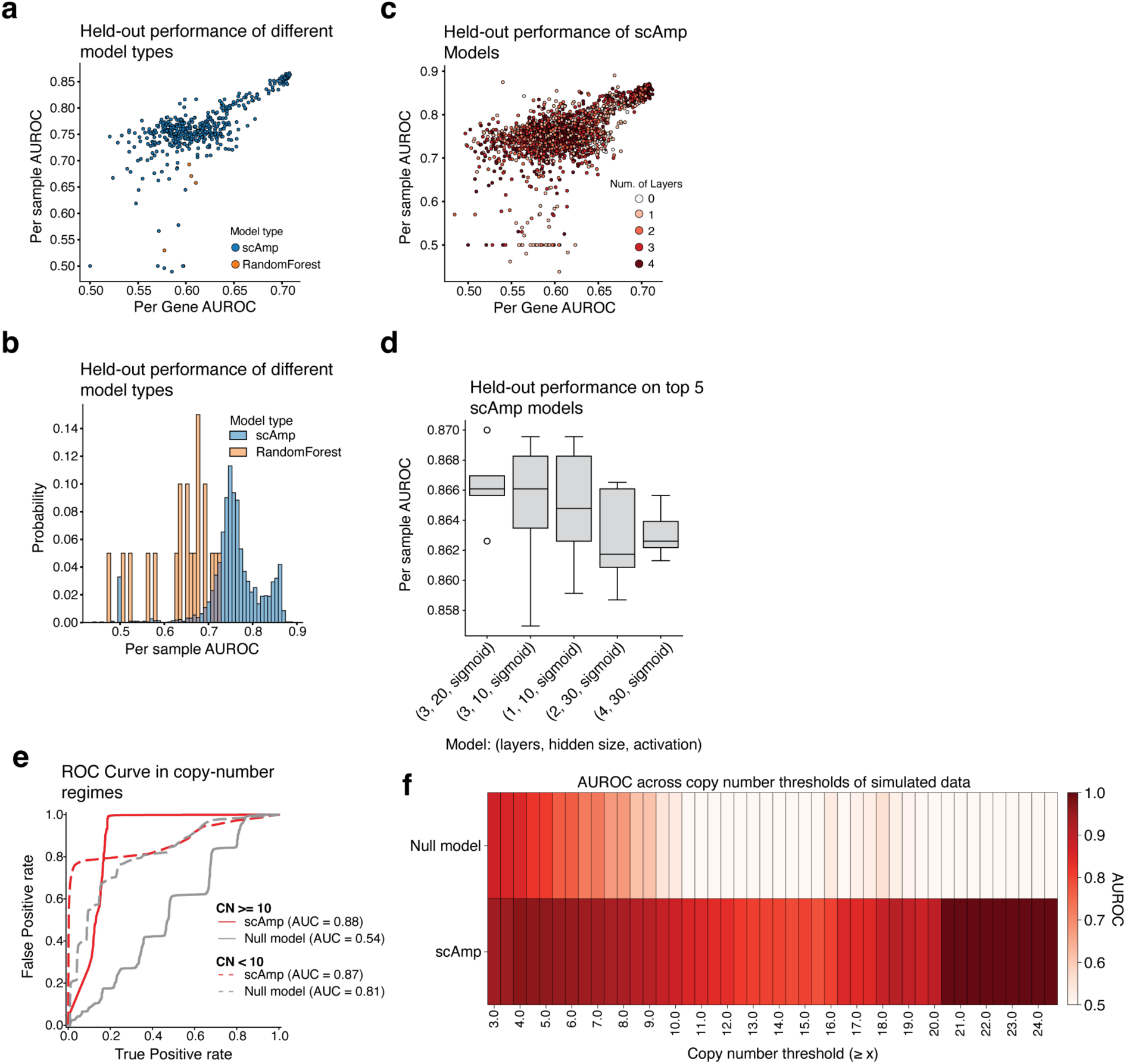
Hyperparameter tuning of *scAmp* models. **(a)** Comparison of performance for all trained *scAmp* and RandomForest classifer models, as quantified by AUROC of per-sample or per-gene ecDNA prediction tasks on held-out patient tumour scATAC-seq data from TCGA. Per-sample ecDNA prediction quantifies the accuracy of predicting whether a patient has at least one ecDNA amplicon; per-gene ecDNA prediction quantifies the accuracy of predicting whether a gene is amplified on ecDNA or not (**Methods**). **(b)** Histogram of AUROC of per-sample ecDNA prediction on held-out patient tumour scATAC-seq data for both RandomForest classifiers and *scAmp* models. **(c)** Scatterplot of *scAmp* model performances colored by number of hidden layers in *scAmp* model. **(d)** Boxplot of the top 5 *scAmp* models, as measured by the AUROC on the per-sample ecDNA prediction task for held-out patient tumour samples. The number of layers, size of hidden layer, and activation function are indicated for each top model. Individual points indicate the performance of a training replicate (i.e., training a model with a new random seed). Boxplots report the quartiles of the distributions, and whiskers extend to 1.5x the interquartile distance. **(e)** ROC curve of *scAmp* and null model on simulated amplicons with average copy-number greater than, or less than, 10. AUROCs are reported in the legend. **(f)** Summary AUROCs of *scAmp* and null model on simulated amplicons subset by a minimum copy-number threshold.

**Extended Data Figure 3.**
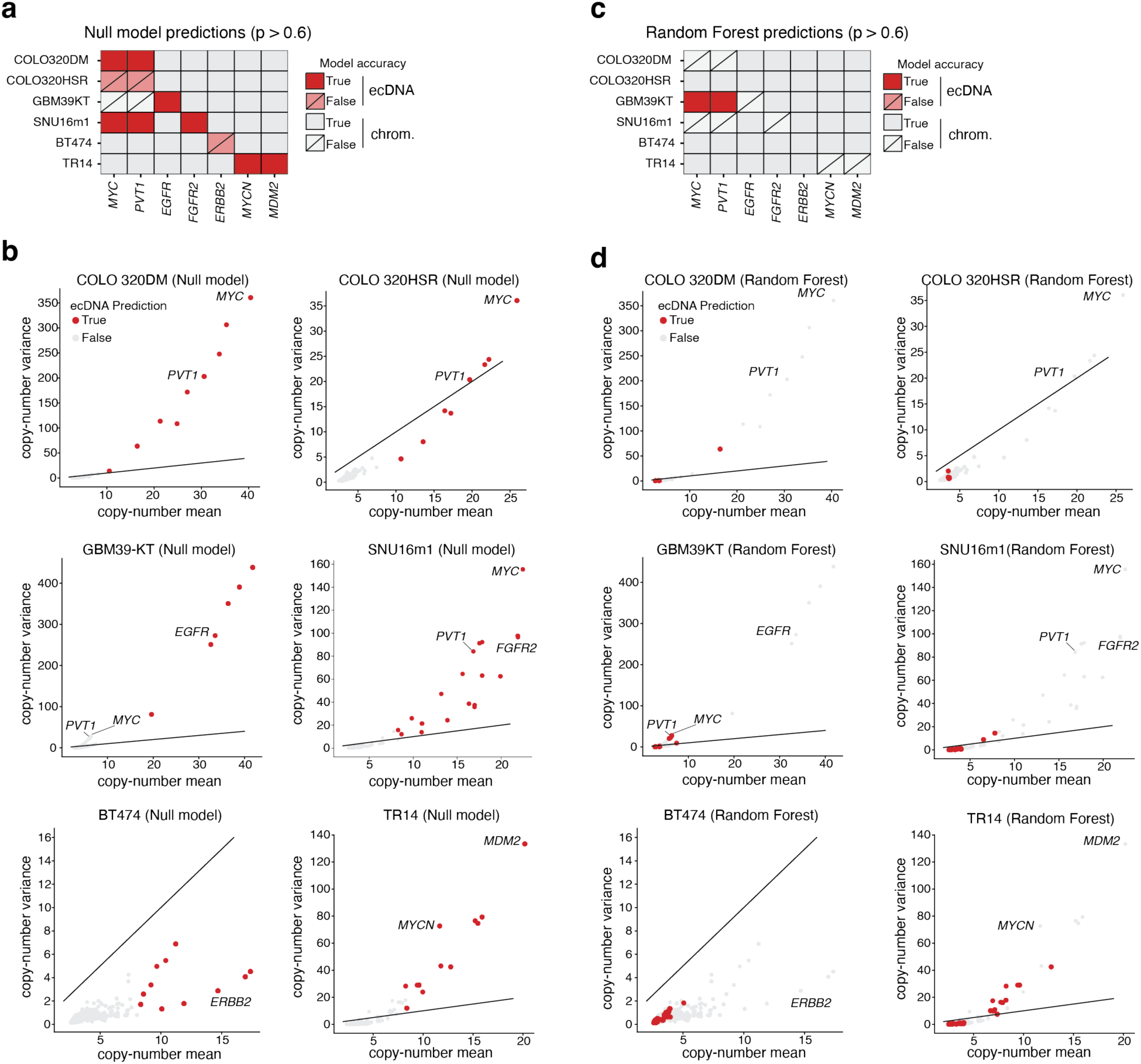
Random Forest and null model performance on held-out cancer cell line data. (a-b) Performance of null linear model (predicting ecDNA status from mean copy-number) on held-out cell line data. **(a)** Summary table of null model predictions of focal genes across cell lines used in this study. Red boxes indicate an ecDNA prediction; grey boxes indicate a chromosomal amplification prediction. Hash marks indicate an incorrect prediction. **(b)** Scatterplots summarizing the mean and variance of gene copy-number distributions and null model predictions in cell lines used in this study. Each point is a single gene, colours indicate model prediction, and focal genes are annotated. **(c-d)** Performance of RandomForest classifier on held-out cell line data. **(c)** Summary table of RandomForest predictions of focal genes across cell lines used in this study. Colours and annotations are the same as in (a). **(d)** Scatterplots summarizing the mean and variance of gene copy-number distributions and RandomForest predictions in cell lines used in this study. Colours and annotations are the same as in (b).

**Extended Data Figure 4.**
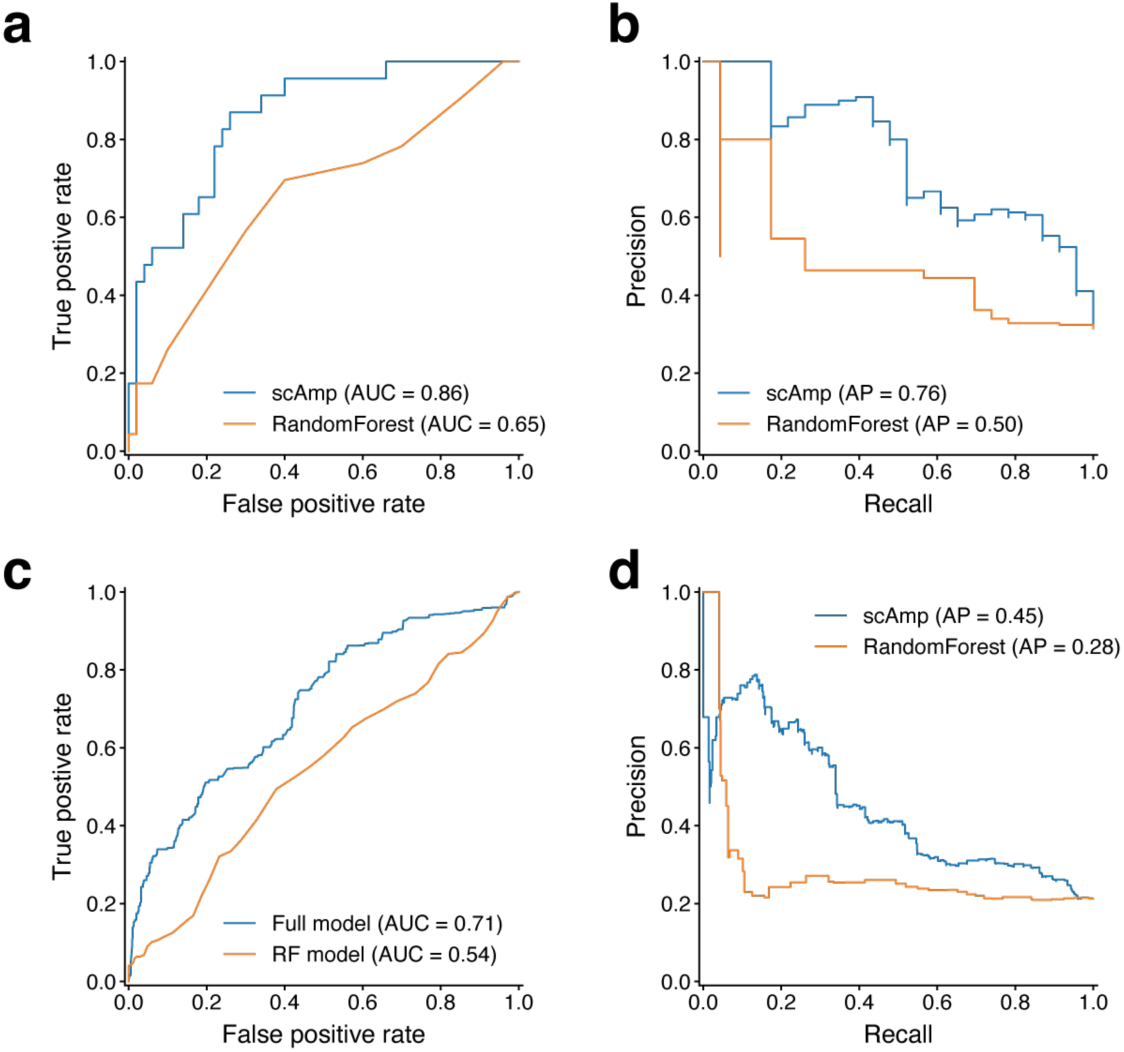
Benchmarking of scAmp 1.0.0 model vs. RandomForest classifier on patient tumour data. A comparison of receiver-operator characteristic (ROC) and precision-recall curves (PRC) for top *scAmp* model and RandomForest classifier on held-out patient tumour scATAC-seq data. **(a-b)** ROC (a) and PRC (b) of models on the per-sample prediction task (predicting whether a given sample has at least one ecDNA+ gene amplification). (**c-d)** ROC (c) and PRC (d) of models on the per-gene prediction task (predicting whether a given gene is amplified on ecDNA in a given tumour sample). AUROC and AUPRCs are reported for each model.

**Extended Data Figure 5.**
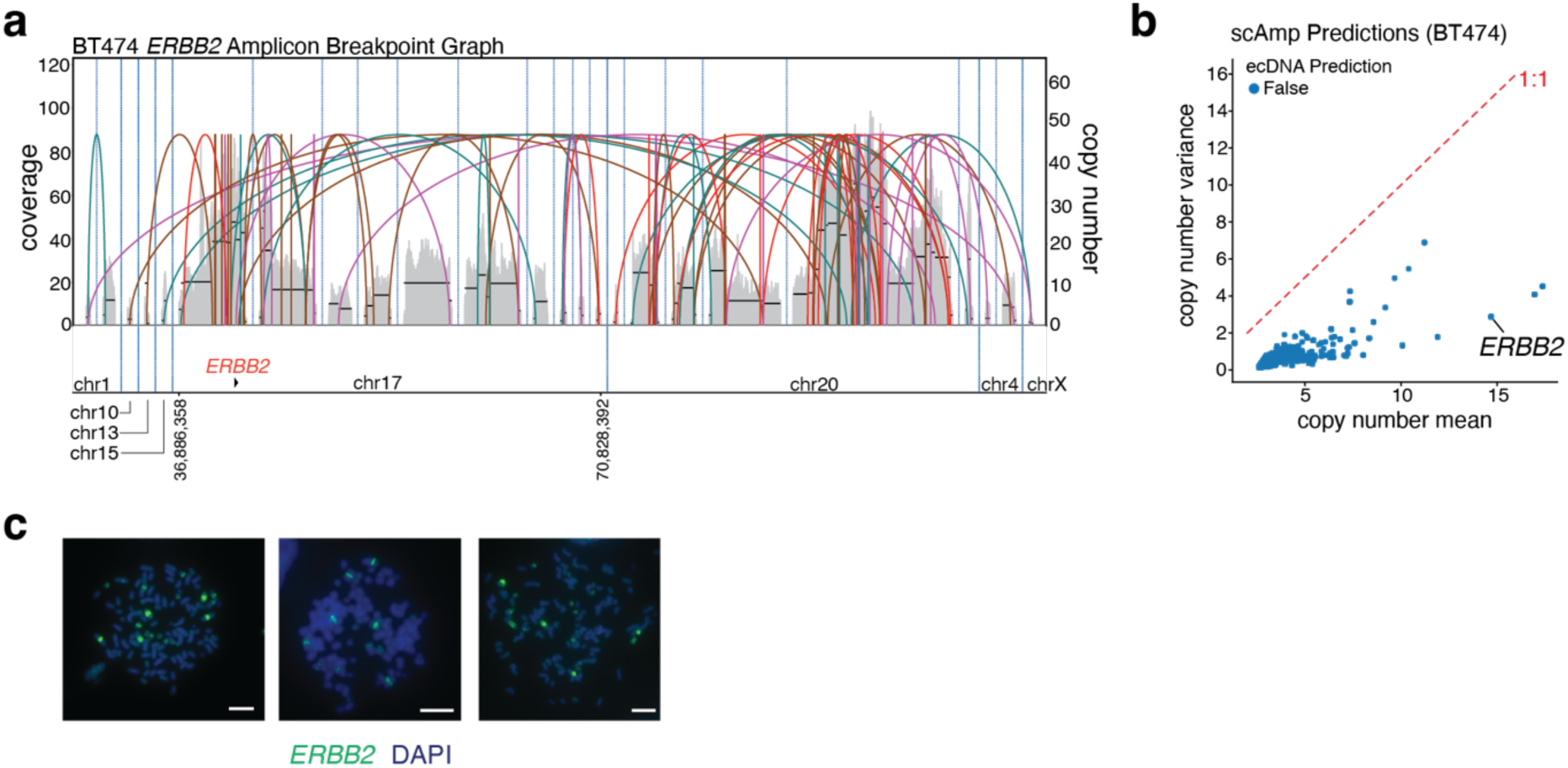
Analysis of *ERBB2* amplification status in the BT474 cell line. **(a)** A breakpoint graph of the ecDNA+ *ERBB2* amplicon produced by AmpliconArchitect^10^ analysis of WGS data (as obtained from AmpliconRepository). The coverage over the amplified intervals is shown as a grey distribution, and each arc represents a breakpoint inferred from AmpliconArchitect. Only the *ERBB2* gene locus is shown underneath for clarity. **(b)** Scatterplots summarizing the mean and variance of gene copy-number distributions for each gene in the BT474 cell line, along with predictions from *scAmp*. *ERBB2* is predicted to be chromosomally amplified. **(c)** Metaphase spread images of cells with DNA FISH for *ERBB2*; here, *ERBB2* co-localizes with DAPI+ chromosomal signal and is inferred to be chromosomally integrated in BT474. Scale bars indicate 10um.

**Extended Data Figure 6.**
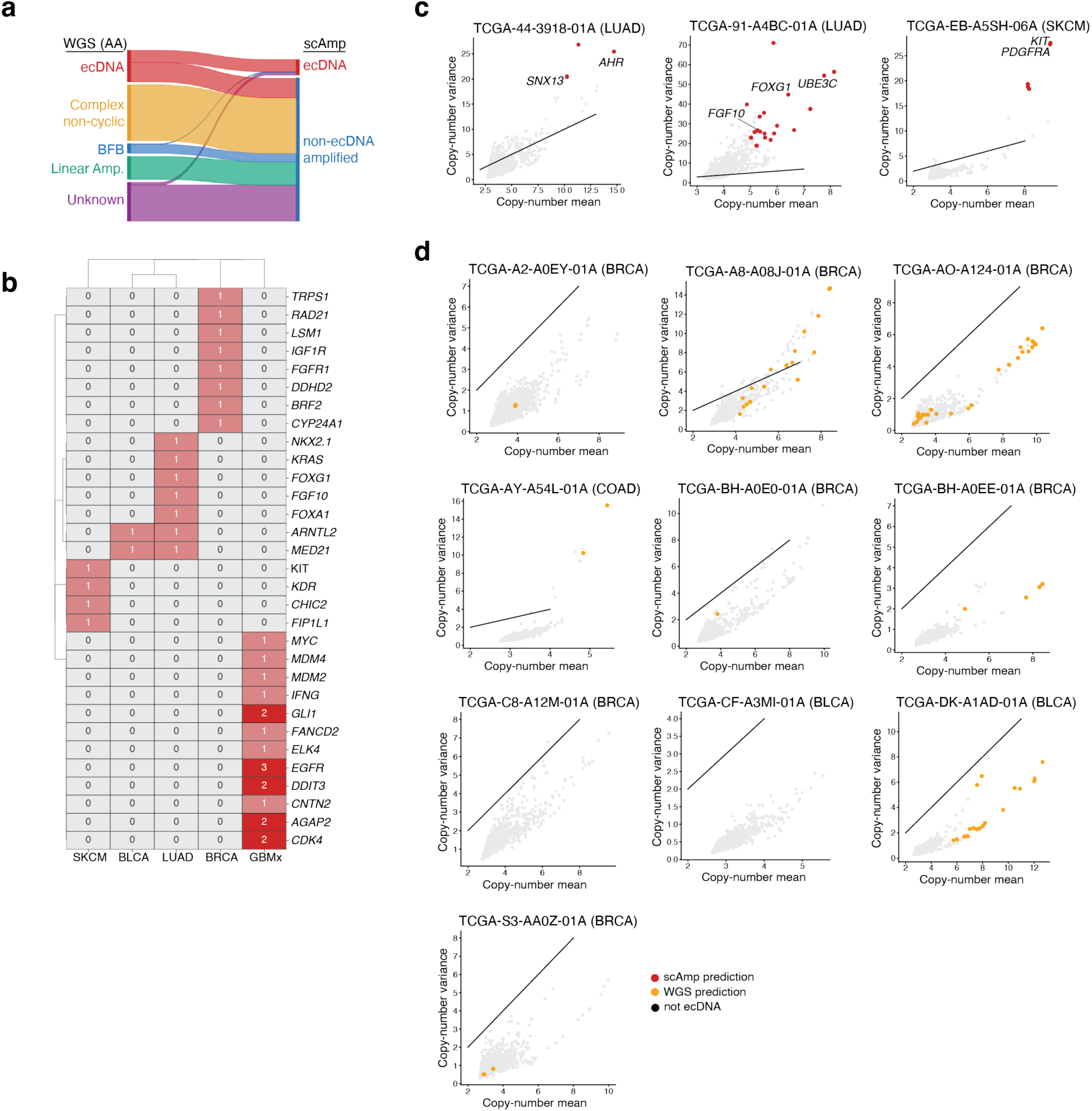
*scAmp* predictions on TCGA scATAC-seq data. **(a)** Comparisons of WGS amplicon predictions versus *scAmp* predictions in the TCGA scATAC-seq cohort. *scAmp* only distinguishes between ecDNA and chromosomal amplifications. Most discrepancies are with WGS-predicted ecDNA amplicons being predicted to be chromosomally amplified by *scAmp*, consistent with analysis of the BT474 cell line. **(b)** Summary of genes most frequently predicted to be amplified on ecDNA by *scAmp* by tumour type. Rows and columns are clustered. **(c-d)** Summary of per-sample discrepancies between *scAmp* and WGS analysis. Scatterplots summarize the mean and variance of gene copy-number distributions for each gene in each patient tumour. Red colours in (c) indicate genes that are predicted by *scAmp* to be ecDNA-amplified, though predicted to be chromosomally amplified by WGS. Orange colours in (d) indicate genes that are predicted by WGS to be ecDNA amplified, though predicted to be chromosomally amplified by *scAmp*. Diagonal lines indicate the 1:1 line. Focal genes of potential interest are called out in (c).

**Extended Data Figure 7.**
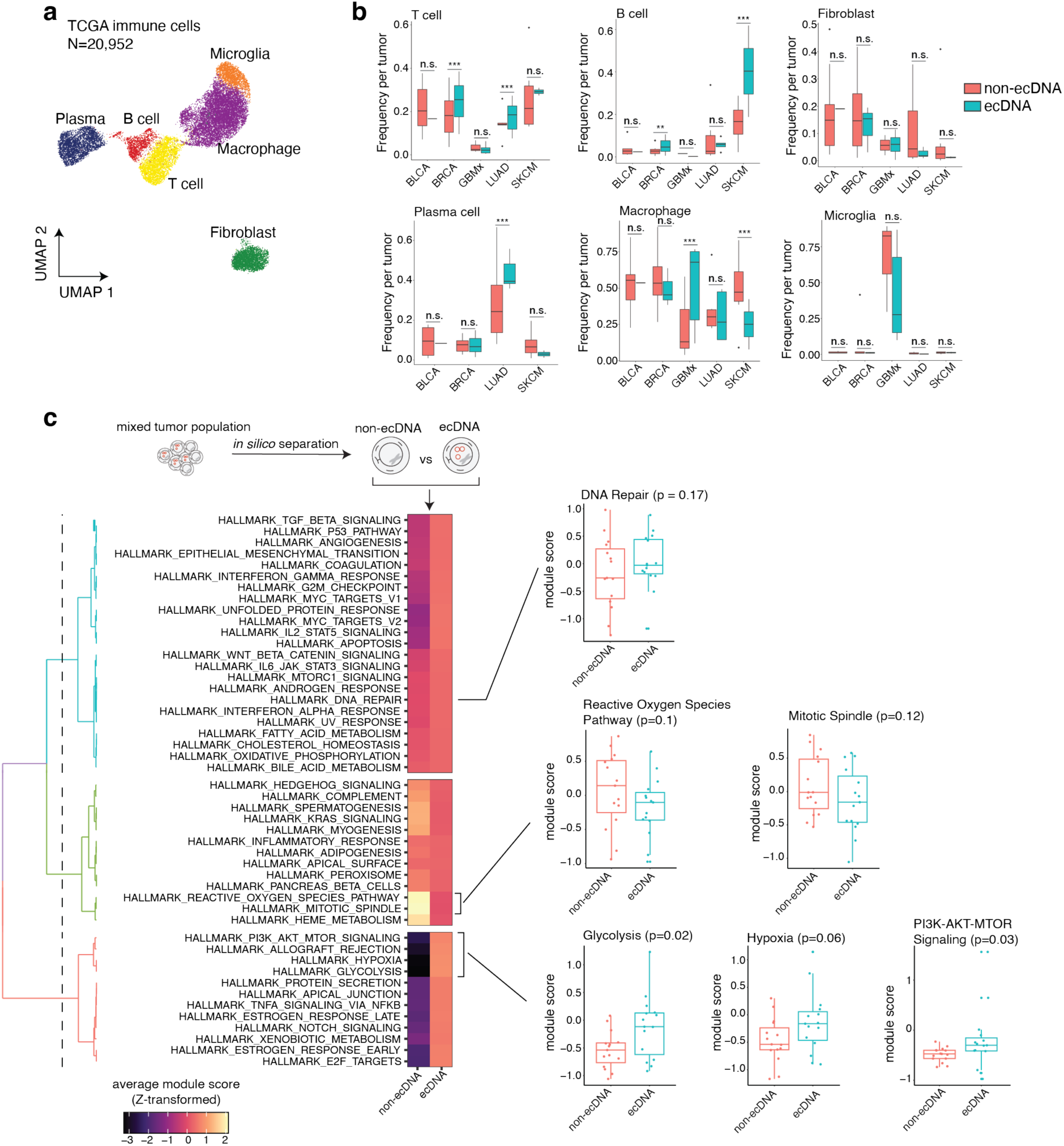
ecDNA-centric analysis of TCGA scATAC-seq data. **(a-b)** Analysis of immune and stromal compartment of TCGA tumors. **(a)** UMAP projection of single-cell chromatin accessibility profiles of immune and stromal cells. Major cell types are annotated. **(b)** Distribution of cell type frequencies in ecDNA+ and non-ecDNA tumours. Frequencies are normalized to the total number of immune and stromal cells profiled in a given tumour (cancer cells excluded). Significances, corresponding to empirical Wilcoxon rank-sums-test *p*-values from 100 random shuffles, are indicated above individual boxplots (***: *p ≤ 0.01*; *\*\*: p ≤ 0.05*). **(c)** Analysis of ecDNA-specific effects on cancer cells. ecDNA+ cancer cells were identified *in silico* based on the copy-number of genes predicted to be on ecDNA (**Methods**). Module scores for Hallmark signature sets were assessed in ecDNA+ and non-ecDNA cells, and average Z-transformed scores are displayed in a clustered heatmap (left). Distributions of individual gene sets are shown in boxplots (right). Each point in the boxplot is the mean module score for the cancer cells in a patient. Boxplots report the quartiles of the distributions, and whiskers extend to 1.5x the interquartile distance. P-values are computed using a one-sided Wilcoxon rank-sums test.

**Extended Data Figure 8.**
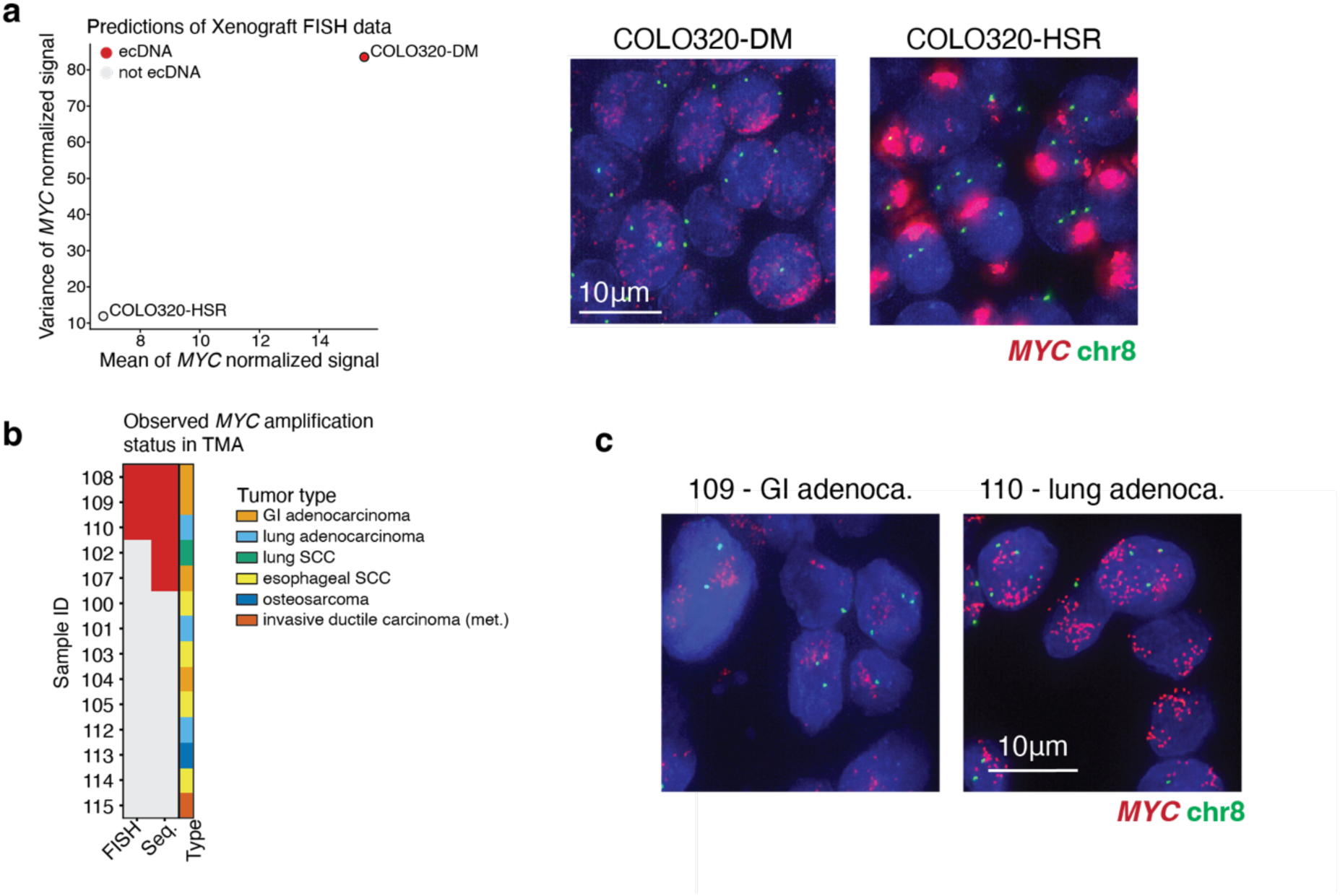
*scAmp* analysis of interphase DNA FISH data. **(a)** Predictions of ecDNA status of *MYC* amplifications in xenograft COLO320-DM and COLO320-HSR tumours. Scatterplots summarize the mean and variance of normalized *MYC* copy-number distributions for COLO320-DM and COLO320-HSR xenograft tumours, as measured with DNA FISH on interphase cells; colours indicate predictions from *scAmp* (left). Representative DNA FISH images from COLO320-DM and COLO320-HSR xenograft tumours; scale bars indicate 10um (right). **(b)** Summary of Tissue Microarray (TMA) obtained of patient tumours; *MYC* amplification status as determined by panel sequencing or DNA FISH is reported for reach sample. Tumour annotations are shown in distinct colours on the right-most bar. **(c)** Representative DNA FISH images for *MYC*-amplified tumour samples; scale-bar indicates 10um. Sample 109 is predicted to be chromosomally amplified by *scAmp* and 110 is predicted to be ecDNA-amplified.

